# Typical visual-field locations facilitate access to awareness for everyday objects

**DOI:** 10.1101/297523

**Authors:** Daniel Kaiser, Radoslaw M. Cichy

**Affiliations:** Department of Education and Psychology, Freie Universität Berlin, Berlin, Germany; Berlin School of Mind and Brain, Humboldt-Universität Berlin, Berlin, Germany; Bernstein Center for Computational Neuroscience Berlin, Berlin, Germany

## Abstract

In real-world vision, humans are constantly confronted with complex environments that contain a multitude of objects. These environments are spatially structured, so that objects have different likelihoods of appearing in specific parts of the visual space. Our massive experience with such positional regularities prompts the hypothesis that the processing of individual objects varies in efficiency across the visual field: when objects are encountered in their typical locations (e.g., we are used to seeing lamps in the upper visual field and carpets in the lower visual field), they should be more efficiently perceived than when they are encountered in atypical locations (e.g., a lamp in the lower visual field and a carpet in the upper visual field). Here, we provide evidence for this hypothesis by showing that typical positioning facilitates an object’s access to awareness. In two continuous flash suppression experiments, objects more efficiently overcame inter-ocular suppression when they were presented in visual-field locations that matched their typical locations in the environment, as compared to non-typical locations. This finding suggests that through extensive experience the visual system has adapted to the statistics of the environment. This adaptation may be particularly useful for rapid object individuation in natural scenes.

## 1. Introduction

Human visual perception is tailored to the world around us: it is most efficient when the input matches commonly experienced patterns. This is evident from low-level vision, where previously experienced regularities determine perceptual interpretations of the input (Purves, Wojtach, & Lotto, 2011). Such influences of typical patterns are also observed for more complex stimuli, such as faces. Face perception is specifically tuned to the typical configuration of facial features (Maurer, Le Grand, & Mondloch, 2001), and a disruption of this configuration (e.g., through face inversion) drastically decreases perceptual performance (Valentine, 1988). Recent studies have suggested that not only the concerted presence of multiple features facilitates face perception, but that also individual facial features profit from typical positioning in the visual field (Chan, Kravitz, Truong, Arizpe, & Baker, 2010, de Haas et al., 2016): for example, it is easier to perceive an eye when it falls into the upper visual field (where it more often appears when looking at a face) than when it falls into the lower visual field (where it is not encountered so often).

Like faces, natural scenes are spatially structured. Scenes consist of arrangements of separable objects, which follow repeatedly experienced configurations (Bar, 2004): for instance, lamps appear above dining tables, and carpets tend to lie on the floor. Previous research has suggested that such typical configurations can facilitate multi-object processing (Draschkow & Võ, 2017; Gronau & Shachar, 2014; Kaiser, Stein, & Peelen, 2014, 2015). It has been proposed that just like in faces, spatial regularities in scenes may also impact the perception of individual objects (Kaiser & Haselhuhn, 2017). As we navigate around, the likelihood of encountering different objects varies across the visual field: for instance, lamps – unless directly fixated – are most often seen in the upper visual field and carpets most often appear in the lower visual field. Because of this repeated expose, typically positioned objects should be processed more efficiently than atypically positioned objects.

To test this hypothesis, we used a variant of continuous flash suppression (CFS; Tsuchiya & Koch, 2005). In breaking-CFS paradigms, a stimulus presented to one eye is temporarily rendered invisible by flashing a dynamic, high contrast mask to the other eye; suppression times, i.e. the time a stimulus needs to break inter-ocular suppression and reach visual awareness, are taken as a measure of processing efficiency (Stein, Hebart, & Sterzer, 2011). Previous studies using this method have shown that suppression times depend on spatial regularity patterns. For example, the typical configuration of faces and bodies facilitates their access to awareness (Jiang, Costello, & He, 2007; Stein, Sterzer, & Peelen, 2012). Similarly, breakthrough is facilitated for typically arranged multi-object configurations (Stein, Kaiser, & Peelen, 2015), demonstrating that the spatial regularities among different objects can facilitate processing under CFS.

To test whether such spatial regularities also impact the processing of individual objects we investigated whether typical retinotopic positioning facilitates an object’s access to awareness. We used a stimulus set consisting of six everyday objects that were either associated with upper or lower visual-field locations (Fig. 1). In two CFS experiments, participants were shown individual exemplars of these objects in their typical or atypical locations onto one eye; a dynamic mask was flashed onto the other eye and temporarily rendered the object invisible (Fig. 2). Participants had to localize the object as fast as possible, irrespective of its identity. In Experiment 1, suppression times (i.e., times until successful localization) were significantly shorter for typically than for atypically positioned objects. In Experiment 2, we replicated this finding, while additionally controlling for potential response conflicts. These results demonstrate that objects appearing in typical visual-field locations gain preferential access to visual awareness, highlighting the influence of natural scene structure on individual object perception.

**Fig. 1.**
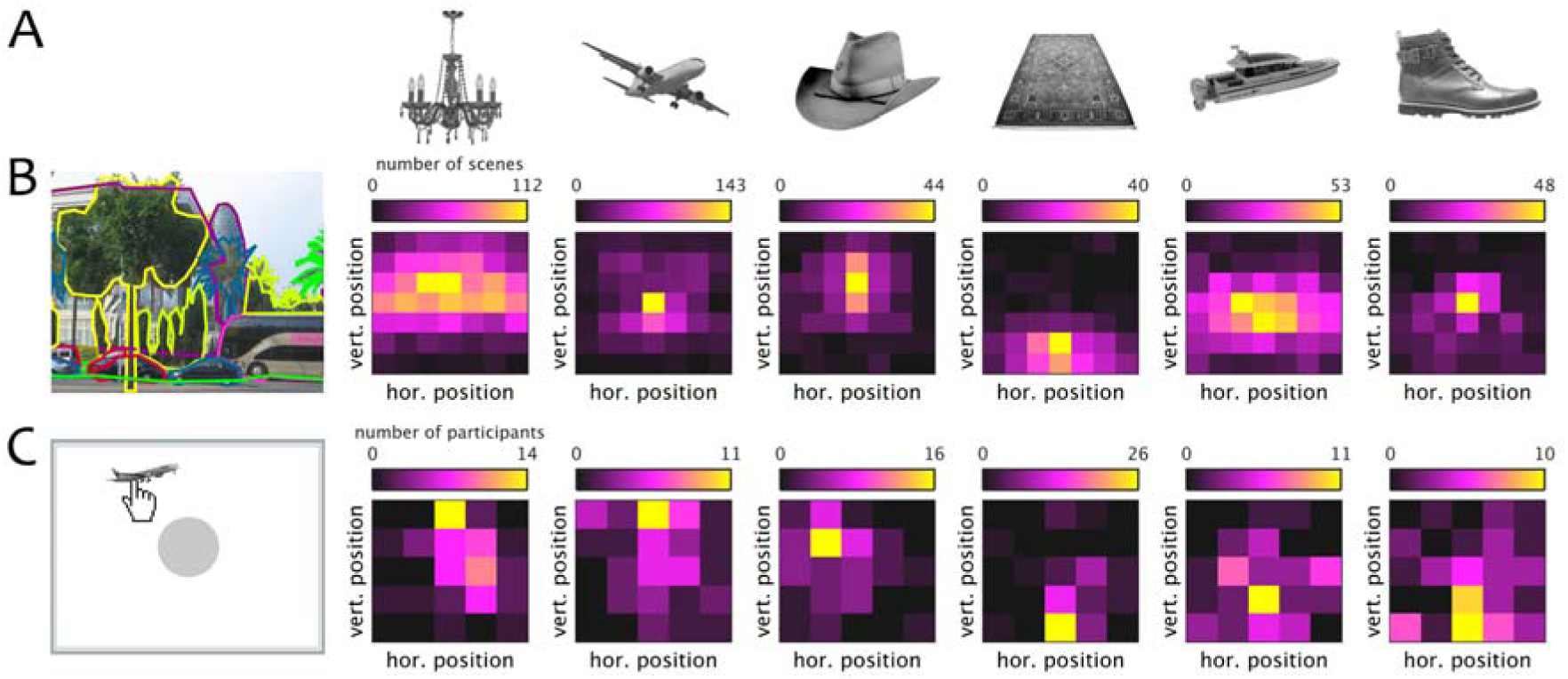
The stimulus set consisted of six objects (10 exemplars each), of which three (lamp, airplane, hat) were associated with upper visual-field locations and three (carpet, boat, shoe) were associated with lower visual-field locations (A). The visual-field associations were validated by computing two measures (see Materials and Methods for details): First, we used a large set of labelled scenes (Russell et al., 2008) to extract typical within-scene positions for each object (B). Second, we asked a set of participants to freely place the object on the screen so that its position best matches its typical real-world position (C). Heatmaps reflect the distribution of locations across a scene (B) or the screen (C).

**Fig. 2.**
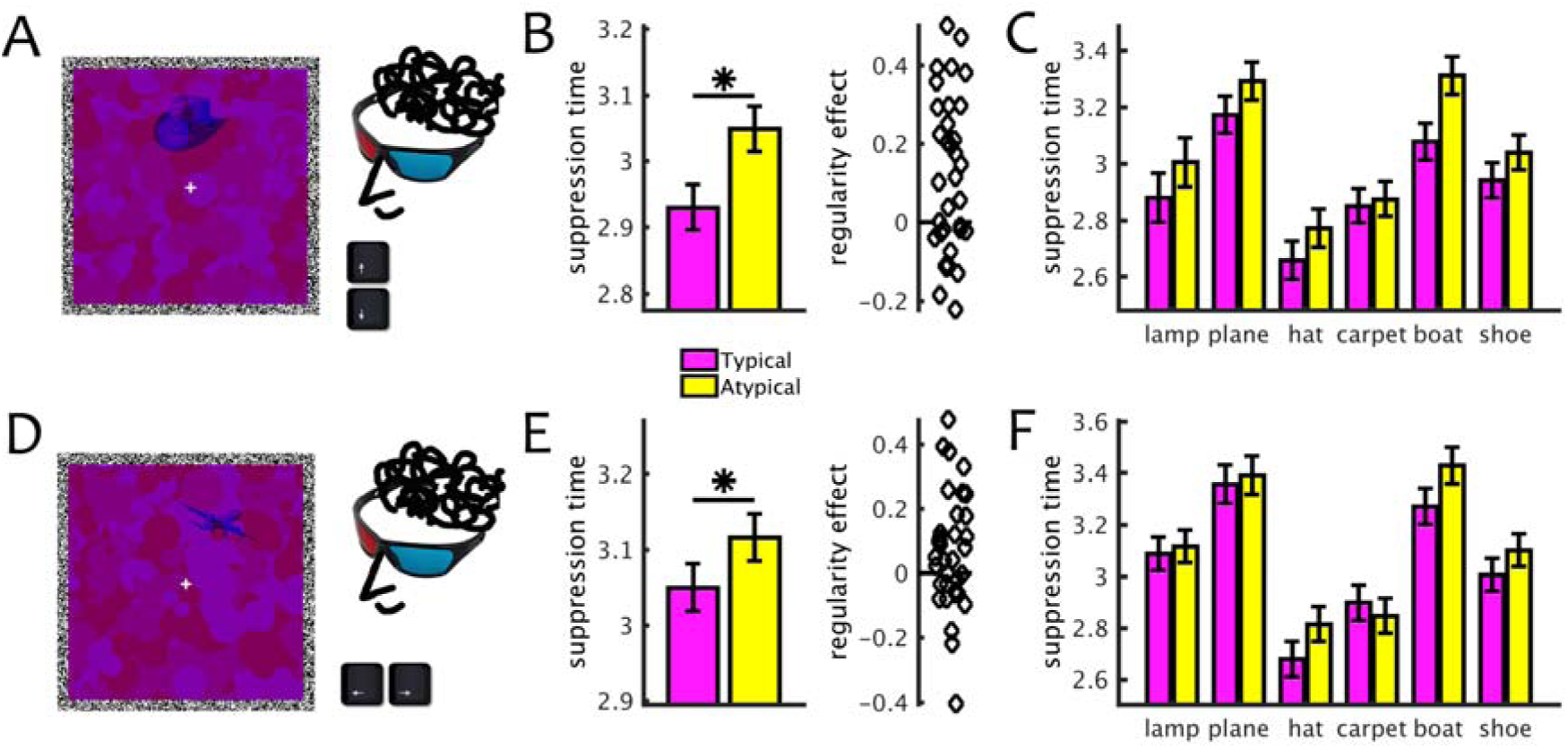
In two CFS Experiments, participants had to localize objects presented to one eye, which were temporarily rendered invisible by dynamic masks presented to the other eye. In Experiment 1, participants had to indicate whether the object appeared in an upper or lower location (A); in Experiment 2, they had to indicate whether it appeared on the left or on the right (D). Crucially, the object could be positioned in its typical location (e.g., hat in the upper visual field) or in an atypical location (e.g., hat in the lower visual field). In both experiments, suppression times were significantly shorter for typically positioned, as compared to atypically positioned, objects (B/E). This effect was numerically consistent across individual objects (but the carpet in Experiment 2) (C/F).

## 2. Material and Methods

### 2.1. Participants

34 healthy adults participated in Experiment 1 (mean age 26.4 years, SD=4.7, 26 female) and another 34 participated in Experiment 2 (mean age 22.9 years, SD=4.4, 26 female). Participants were recruited from the online participant database of the Berlin School of Mind and Brain (Greiner, 2005). All participants had normal or corrected-to-normal vision, provided informed consent and received monetary reimbursement or course credits for participation. All procedures were approved by the local ethical committee and were in accordance with the Declaration of Helsinki.

Sample size was determined by an a-priori power calculation: assuming a hypothetical, medium-sized effect of d=0.5, 34 participants are needed for a power of 80%^1^.

### 2.2. Stimuli

The stimulus set consisted of six objects (Fig. 1A). Three of the objects were associated with upper visual-field locations (lamp, airplane, and hat) and three were associated with lower visual-field locations (carpet, boat, and shoe). For each object, we collected ten exemplars. The objects were matched for their categorical content (two furniture items, two transportation items, and two clothing items) to match high-level properties (e.g., the objects’ size, manipulability and semantic associations) across upper and lower visual-field objects. To control for low-level confounds, stimulus images were gray-scaled and matched for overall luminance (Willenbockel et al., 2010). Additionally, we checked whether there was a consistent low-level difference across objects associated with upper and lower visual-field objects. For this, we computed pair-wise pixel correlations for all conditions, and compared results for objects associated with the same visual-field locations versus objects associated with different visual-field locations. This test was not significant, t(1498)=0.50, p=0.62, suggesting that there was no consistent low-level difference across upper and lower visual-field objects.

To validate the objects’ associations with specific locations, we used two complementary approaches. First, we automatically queried a large database (>10,000 images) of labelled scene photographs (LabelMe; Russell, Torralba, Murphy, & Freeman, 2008). We assumed that the distribution of objects across a larger number of photographs approximates their distribution under natural viewing conditions. For each scene that contained one of the six objects, we extracted the within-scene location (the mean coordinate of the labelled area) of the object (Fig. 1B). Second, we explicitly asked a set of participants to place each object on a computer screen such that its on-screen position mirrored its most probable real-world positioning (Fig. 1C). For both validation approaches, vertical locations were significantly higher for upper than for lower visual-field objects (all t>6.04, p<.001). Both measures thus confirmed the objects’ associations with specific, typical locations. A detailed report of our validation procedure can be found in Kaiser, Moeskops, and Cichy (2018).

### 2.3. Experimental Design

The design was identical for both CFS experiments, unless otherwise noted. During the experiment, participants wore red/blue anaglyph glasses, which allowed for a separation of the two eye channels. Each stimulus display consequently consisted of a combination of red and blue stimulus layers: One layer (“stimulus layer”) contained the object stimulus, while the other layer (“mask layer”) contained a flashing noise mask.

The stimulus layer contained one exemplar of one of the six objects, shown on a uniform-intensity background. In Experiment 1, the object (max. 3° visual angle) could appear in one of two locations (3° eccentricity), either in the upper or the lower visual field (Fig. 2A). In Experiment 2, the objects appeared in one of four locations, where the upper and lower locations were additionally shifted either to the right or to the left (by 1.5° visual angle) (Fig. 2D). The stimulus layer was always presented to the participant’s non-dominant eye.

The mask layer contained dynamic, contour-rich CFS masks consisting of randomly arranged white, black, and gray circles (see Figure 2A/D). These masks were re-drawn every 100ms, so that the mask layer flickered at a frequency of 10Hz. The mask layer was always presented to the participant’s dominant eye.

During each trial, the stimulus display appeared within a square frame (12° visual angle width/height, consisting of a black-and-white noise contour), placed on a black background. In the center of the frame, a white fixation cross was overlaid onto the stimulus; participants were instructed to maintain central fixation throughout the experiment. To avoid abrupt gradients, the stimulus layer was gradually faded in over the first second of each trial (by linearly increasing its contrast) and then remained constant until the end of the trial. If participants had not responded after eight seconds, the mask layer was faded out over the next four seconds (by linearly decreasing its contrast). Participants had to indicate in which part of the screen they saw an object by using the arrow keys on the keyboard. In Experiment 1, participants had to indicate whether the object appeared in the upper or lower position within the box (Fig. 2A). In Experiment 2, participants had to indicate whether the object appeared to the right or the left of the vertical midline (Fig. 2D). In both experiments, participants were instructed to respond as fast as possible when any part of the target stimulus became visible, irrespectively of their recognition of the object. Trials were terminated as soon as participants responded, followed by an inter-trial interval of one second.

Before the start of the experiment, participants completed a short familiarization block (around 5 minutes, containing a random subset of experimental trials). After this familiarization block, mask contrast was adjusted for some participants, to avoid very short or very long breakthrough times. Importantly, within participants, the mask contrast remained identical for all trials of the subsequent experiment. Both experiments contained 480 trials. In Experiment 1, each object exemplar appeared four times in each of the two locations. In Experiment 2, each object exemplar appeared two times in each of the four locations. Trial order was fully randomized. Participants could take breaks after 120, 240, and 360 trials. Stimulus presentation was controlled using the Psychtoolbox (Brainard, 1997).

### 2.4. Statistical analysis

Trials with wrong responses or suppression times <300ms were discarded from all analysis. Suppression times were then averaged by typicality, i.e. separately for typically and atypically positioned objects. Statistical significance was assessed using paired t-tests^2^. Cohen’s d is reported as an effect-size measure for t-tests.

For computing object-specific regularity effects, we first corrected for bias towards either the upper or lower visual field in individual participants’ responses (e.g., caused by preferences in attentional allocation). To account for these biases in individual participants, we first computed the suppression time difference between objects appearing in the upper and lower locations (independently of positional regularities). Then, we subtracted away half of this difference from all suppression times for the “slower” location, and added half of this difference to all suppression times for the “faster” location. Effects were then compared across objects using repeated-measures ANOVAs. Partial *ƞ*^2^ is reported as an effect-size measure for ANOVAs.

## 3.1 Results

In both experiments, participants localized the objects with high accuracy (Experiment 1: 99%; Experiment 2: 98%). Accuracy was not analyzed further.

### 3.1. Experiment 1

In Experiment 1, we tested whether typical visual-field locations facilitate object perception under inter-ocular suppression. Participants had to indicate as fast as possible whether the object appeared above or below fixation (Fig. 2A). Crucially, suppression times were significantly shorter for typically positioned objects (e.g., a hat in the upper visual field) than atypically positioned objects (e.g., a hat in the lower visual field), t(33)=3.45, p=.002, d=0.59 (Fig. 2B), suggesting that typical object positioning boosts access to visual awareness.

### 3.2. Experiment 2

In Experiment 2, we replicated the findings obtained in Experiment 1. We additionally sought to exclude potential response conflicts (e.g., a “down” motor response could, in principle, conflict with the object having a typical “upper” location); we therefore asked participants to indicate whether the object appeared shifted to the right or left of the vertical midline (Fig. 2D). Again, suppression times were shorter for typically positioned objects, t(33)=2.12, p=.042, d=0.36 (Fig. 2E), corroborating the finding that typical object locations facilitate access to awareness.

### 3.3. Individual-object effects

To compare the regularity benefit across objects, we examined suppression times for individual objects when they were positioned typically or atypically (see Materials and Methods). Notably, a net facilitation of detection was found for each object in Experiment 1 (Fig. 2C), and for all but one objects (carpet) in Experiment 2 (Fig. 2F). In both experiments, no modulation of this regularity benefit was found across individual objects, Experiment 1: F(5,165)=1.04, p=.40, 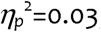, Experiment 2: F(5,165)=0.37, p=.87, 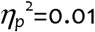. This pattern of results demonstrates that the effects were consistent across objects and not driven by individual stimuli.

## 4. Discussion

Here, we provide evidence that typical visual-field locations facilitate the perception of everyday objects under inter-ocular suppression. In two CFS experiments, objects appearing in their typical visual-field locations had shorter suppression times than objects appearing in atypical locations. In both experiments, this benefit was consistent across individual objects. Experiment 2 additionally ruled out response conflicts as an alternative explanation for the effect. By showing that conjunctions of objects and locations are differentially likely to enter visual awareness, our findings highlight the impact of real-world statistics on perceptual processing.

Our results complement recent studies showing that breakthrough under CFS is modulated by regularities in multi-object arrangements (Stein et al., 2015) and by congruencies between objects and their scene context (Mudrik, Breska, Lamy, & Deouell, 2011; but see Moors, Boelens, van Overwalle, & Wagemans, 2016). Together, this indicates that the human visual system is tuned to spatial regularities at different levels of complexity – from regularities in individual object positioning to spatial dependencies within a complex scene. In real-life situations, tuning to these different spatial regularities may offer complementary contributions to efficient object perception in complex, but regularly organized, environments (Wolfe, Võ, Evans, & Greene, 2011). Interestingly, the results from CFS paradigms suggest that the presence of regularities may not only facilitate conscious and explicit interactions with the world, but may also determine whether we perceive an object in the first place.

What allows typically positioned objects to overcome inter-ocular suppression more efficiently? There is considerable agreement that processing under inter-ocular suppression is unlikely to suffice for a full semantic analysis (Gayet, Van der Stigchel, & Paffen, 2014; Lin & He, 2009). However, numerous studies have demonstrated that processing under CFS is modulated by experience: for example, access to awareness is facilitated for familiar faces (Gobbini et al., 2013), own-race faces (Stein, End, & Sterzer, 2014), objects of expertise (Stein, Reeder, & Peelen, 2016), and typically arranged multi-object arrangements (Stein et al., 2015). Our results similarly reflect a benefit of extensive experience, induced by life-long exposure to particular object-location conjunctions.

On a neural level, extensive experience can sharpen tuning properties in visual cortex (Freedman, Riesenhuber, Poggio, & Miller, 2006; Kobatake, Wang, & Tanaka, 1998). Two recent fMRI studies have suggested that typically positioned face parts evoke more distinctive response patterns, suggesting that typically experienced retinotopic locations sharpen visual representations for facial features (Chan et al., 2010; de Haas et al., 2016). Similar results have been obtained in an EEG experiment using the same stimuli as the current study (Kaiser et al., 2018), providing evidence for more distinctive cortical representations than for typically positioned objects. Interestingly, it has been suggested that the ability of an object to overcome inter-ocular suppression is tied to the distinctiveness of its neural representation (Cohen, Nakayama, Konkle, Stantic, & Alvarez, 2015). The sharpened cortical representations for typically positioned objects might thus form a cortical correlate of the objects’ increased ability to overcome inter-ocular suppression.

To conclude, our findings reveal how spatial regularities in natural environments impact perceptual processing of individual objects: when objects appear in typical locations, their access to visual awareness is facilitated. This facilitation may be a valuable prerequisite for fast object individuation in complex real-world scenes.

## Acknowledgements

We thank Merle Moeskops for her help with stimulus preparation and the collection of the behavioral validation data. The research was supported by a DFG Emmy Noether Grant awarded to R.M.C. (CI241-1/1).

## Footnotes

1 A power analysis based on the effect obtained in Experiment 1 (d=0.59) revealed a power of 92% for a sample size of 34 in Experiment 2.

2 In both experiments, differences in suppression times were approximately normally distributed (Shapiro-Wilk tests: both W>0.96, p>.27).

## References

Bar, M. (2004). Visual objects in context. Nature Reviews Neuroscience, 5, 617–629.

Brainard, D. H. (1997). The psychophysics toolbox. Spatial Vision, 10, 433–436.

Chan, A. W., Kravitz, D. J., Truong, S., Arizpe, J., & Baker, C.I. (2010). Cortical representations of bodies and faces are strongest in commonly experienced configurations. Nature Neuroscience, 13, 417–418.

Cohen, M. A., Nakayama, K., Konkle, T., Stantic, M., & Alvarez, G. A. (2015). Visual awareness is limited by the representational architecture of the visual system. Journal of Cognitive Neuroscience, 27, 2240–2252.

de Haas, B., Schwarzkopf, D. S., Alvarez, I., Lawson, R. P., Henriksson, L., Kriegeskorte, N., & Rees, G. (2016). Perception and processing of faces in the human brain is tuned to typical facial feature locations. Journal of Neuroscience, 36, 9289–9302.

Draschkow, D., & Võ, M. L.-H. (2017). Scene grammar shapes the way we interact with object, strengthens memories, and speeds search. Scientific Reports, 7, 16471.

Freedman, D. J., Riesenhuber, M., Poggio, T., & Miller, E. K. (2006). Experience-dependent sharpening of visual shape selectivity in inferior temporal cortex. Cerebral Cortex, 16, 1631–1644.

Gayet, S., Van der Stigchel, S., & Paffen, C. L. (2014). Breaking continuous flash suppression: competing for consciousness on the pre-semantic battlefield. Frontiers in Psychology, 5, 460.

Gobbini, M. I., Gors, J. D., Halchenko, Y. O., Rogers, C., Guntupalli, J. S., Hughes, H., & Cipolli, C. (2013). Prioritized detection of personally familiar faces. PLoS One, 8, e66620.

Greiner, B. (2015). Subject Pool Recruitment Procedures: Organizing Experiments with ORSEE. Journal of the Economic Science Association, 1, 114–125.

Gronau, N., & Shachar, M. (2014). Contextual integration of visual objects necessitates attention. Attention, Perception & Psychophysics, 76, 695–714.

Jiang, Y., Costello, P., & He, S. (2007). Processing of invisible stimuli: advantage of upright faces and recognizable words in overcoming interocular suppression. Psychological Science, 18, 349–355.

Kaiser, D., & Haselhuhn, T. (2017). Facing a regular world: How spatial object structure shapes visual processing. Journal of Neuroscience, 37, 1965–1967.

Kaiser, D., Moeskops, M. M., & Cichy, R. M. (2018). Typical real-world locations impact the time course of object coding. bioRxiv, https://doi.org/10.1101/177493.

Kaiser, D., Stein, T., & Peelen, M. V. (2014). Object grouping based on real-world regularities facilitates perception by reducing competitive interactions in visual cortex. Proceedings of the National Academy of Sciences, U.S.A., 111, 11217–11222.

Kaiser, D., Stein, T., & Peelen, M. V. (2015). Real-world spatial regularities affect visual working memory for objects. Psychonomic Bulletin & Review, 22, 1784–1790.

Kobatake, E., Wang, G., & Tanaka, K. (1998). Effects of shape-discrimination training on the selectivity of inferotemporal cells in adult monkeys. Journal of Neurophysiology, 80, 324–330.

Kravitz, D. J., Vinson, L. D., & Baker, C. I. (2008). How position dependent is visual object recognition? Trends in Cognitive Sciences, 12, 114–122.

Lin, Z., & He, S. (2009). Seeing the invisible: the scope and limits of unconscious processing in binocular rivalry. Progress in Neurobiology, 87, 195–211.

Maurer, D., Le Grand, R., & Mondloch, C. J. (2001). The many faces of configural processing. Trends in Cognitive Sciences, 6, 255–260.

Moors, P., Boelens, D., van Overwalle, J., & Wagemans, J. (2016). Scene integration without awareness: no conclusive evidence for processing scene congruency during continuous flash suppression. Psychological Science, 27, 945–956.

Mudrik, L., Breska, A., Lamy, D., & Deouell, L. Y. (2011). Integration without awareness: expanding the limits of unconscious processing. Psychological Science, 22, 764–770.

Purves, D., Wojtach, W. T., & Lotto, R. B. (2011). Understanding vision in wholly empirical terms. Proceedings of the National Academy of Sciences, U.S.A., 108, 15588–15595.

Russell, B. C., Torralba, A., Murphy, K. P., & Freeman, W. T. (2008). LabelMe: a database and web-based tool for image annotation. International Journal of Computer Vision, 77, 157–173.

Stein, T., End, A., & Sterzer, P. (2014). Own-race and own-age biases facilitate visual awareness of faces under interocular suppression. Frontiers in Human Neuroscience, 8, 582.

Stein, T., Hebart, M. N., & Sterzer, P. (2011). Breaking continuous flash suppression: A new measure of unconscious processing during interocular suppression? Frontiers in Human Neurosciences, 5, 167.

Stein, T., Kaiser, D., & Peelen, M. V. (2015). Interobject grouping facilitates visual awareness. Journal of Vision, 15, 10.

Stein, T., Reeder, R. R., & Peelen, M. V. (2016). Privileged access to awareness for faces and objects of expertise. Journal of Experimental Psychology: Human Perception & Performance, 42, 788–798.

Stein, T., Sterzer, P., & Peelen, M. V. (2012). Privileged detection of conspecifics: evidence from inversion effects during continuous flash suppression. Cognition, 125, 54–79.

Tsuchiya, N., & Koch, C. (2005). Continuous flash suppression reduces negative afterimages. Nature Neuroscience, 8, 1096–1101.

Valentine, T. (1988). Upside-down faces: A review of the effect of inversion upon face recognition. British Journal of Psychology, 79, 471–491.

Willenbockel, V., Sadr, J., Fiset, D., Horne, G. O., Gosselin, F., & Tanaka, J. W. (2010). Controlling low-level image properties: The SHINE toolbox. Behavior Research Methods, 42, 671–684.

Wolfe, J. M., Võ, M. L.-H., Evans, K. K., & Greene, M. R. (2011). Visual search in scenes involves selective and nonselective pathways. Trends in Cognitive Sciences, 15, 77–84.

